# 4-week GLP immunotoxicity assessment of lactoferrin alpha produced by *Komagataella phaffii* in Sprague Dawley rats

**DOI:** 10.1101/2024.08.16.608335

**Authors:** Ross Peterson, Robert B. Crawford, Lance K. Blevins, Norbert E. Kaminski, Anthony J. Clark, Carrie-Anne Malinczak

## Abstract

Oral toxicity and toxicokinetic properties of human lactoferrin (LF) alpha produced in *Komagataella Phaffii* (effera™) were investigated in adult Sprague-Dawley rats over a 28-day period under good laboratory practice conditions. Main study dosing used groups of 10 rats/sex/dose, and a secondary study evaluating toxicokinetic parameters used 6 rats/sex/dose. The vehicle control group received sodium citrate buffer, test groups received daily doses of 200, 600, and 2000 mg of effera™ per kg body weight, and the comparative control group received 2000 mg bovine LF (bLF)/kg body weight per day. T-cell dependent antibody response against keyhole limpet hemocyanin and immunophenotyping of the spleen were performed as measures of immunotoxicity. Clinical observations, body weight, hematology, coagulation, clinical chemistry, urinalysis, immunotoxicity, gross necropsy, and histopathology were assessed. Toxicokinetic parameters were analyzed as an indication of LF bioavailability, and anti-LF antibody assays were conducted to detect antibodies produced against LF to measure immunogenicity. No treatment related toxicologically significant changes were observed. Based on the absence of toxicologically relevant changes, effera™ is well tolerated in rats at doses up to 2000 mg rhLF/kg/day, an amount ∼400 times that of the estimated daily intake at the 90th percentile proposed for human adult use.

## INTRODUCTION

Lactoferrin (LF) is an 80 kDa member of the transferrin family of glycoproteins, produced by exocrine glands (e.g., mammalian milk or tears) and by neutrophils. It is a nutrient, primarily involved in host immunity and iron homeostasis.^1,2^ It is known that several of the functional attributes of lactoferrin are explained by its capacity to bind iron and behave as a mediator of the innate and adaptive immune systems.

Human LF (hLF) and bovine LF (bLF) are similar in some respects yet have notable differences in terms of their amino acid identity and glycosylation patterns that may impact their biological functions. The first bLF was isolated in 1939 by Sorenson and Sorenson,^3^ while hLF was later isolated from human milk (hmLF) in the 1960s by Johansson.^4^ Both hLF and bLF have approximately 691 and 700 amino acid residues, respectively. Bovine LF is Generally Recognized as Safe (GRAS) for use in numerous conventional foods as well as in infant formula and has various applications in the food industry.^5–8^ Physiological differences in functionality between bLF and hLF suggest that human forms of LF have the potential to be more functionally active than bLF as a food ingredient. For example, hLF has been shown to digest more slowly, has stronger affinity for the human LF receptor in the small intestine, and has a different digestive profile following human enzyme digestion, and therefore may be more bioavailable than bLF.^9^ Additionally, a recent study showed that different forms of hLF are similar to each other but divergent from bLF upon digestion and bioactive peptide release,^10^ supporting closer functionality between the human forms. Finally, the human form of LF, versus bLF, may be less immunogenic given its identical native amino acid sequence.

Several groups have used transgenic expression systems to produce a recombinant form of hLF (rhLF) to circumvent ethical and cost constraints present with mass production of native hLF from human milk. While the amino acid profile can be identical between native hLF and rhLF, their profiles may differ due to differences in their N-glycosylation patterns, presence of residual host proteins in rhLF, and other distinctive characteristics. Expression systems to produce rhLF have included fungi, rice, transgenic cattle, and yeast, and there have been multiple toxicology studies conducted using these products. For example, Cerven et al. found a lack of general toxicity of rice-derived human apo-lactoferrin (low iron saturation) in a 28-day study in Wistar rats,^11^ and a similar study by the same group also found no toxic effects of human holo-lactoferrin (high iron saturation) in Wistar rats.^12^ Appel et al. found no toxicologically relevant results in a 13-week oral toxicity assessment in Wistar rats with transgenic cattle-derived rhLF.^13^ A similar study conducted in Sprague Dawley rats fed a diet for 90 days consisting of up to 30% rhLF-containing transgenic milk powder also found no adverse effects on clinical, biological, and pathological parameters and concluded that the milk powder product containing rhLF is as safe as conventional milk.^14^

Helaina Inc. (New York City, New York, USA) successfully developed lactoferrin alpha (effera™) which has the same amino acid profile as hLF and is produced in a glycoengineered yeast.^15^ Briefly, a modified yeast strain of *Komagataella phaffii* uses a proprietary technology where the endogenous glycosyltransferase gene (OCH1) is disrupted and a stepwise introduction of heterologous glycosylation enzymes enables *K. phaffii* to produce hLF. The glycoprotein is then purified using a proprietary process and contains no detectable strain DNA down to 10 femtograms, and no antibiotic-resistant genes. Full identity characteristics of effera™ were recently published and found that effera™ is similar in identity to that of native hLF controls, with the exception of N-glycan groups and iron saturation.^15^ Native LF is produced ubiquitously in the human body with exposure present in biological fluids and mucous secretions. The concentration of hLF is highest in hmLF, from 5–7 mg/mL in colostrum to 1–3 mg/mL in mature milk therefore, breastfed babies are exposed to high levels of hmLF in early life.^16,17^ Additionally, adults are also constantly exposed to hLF naturally occurring in saliva (7-9 mg/L),^18^ tears (∼0.62 g/L)^19^ and other epithelial cell secretions.^20^ Therefore, oral consumption of effera™ is not anticipated to yield toxicological effects given its similarities to the native form.

However, despite these similarities, safety assessments are critical for any food ingredient being produced by a novel production organism.^21^ In addition to general toxicological analysis, given well-known immunomodulatory properties of hLF, immunotoxicity potential should also be evaluated. Few previous studies have examined changes in immune function by rhLF that provide insights into its putative immunotoxicologic effects, crucial for ensuring safety. A previous study conducted using effera™ was the first of its kind to include a comprehensive evaluation of its effects on the frequency of major immune cell types by immunophenotyping.^22^ Immunophenotyping is an application of flow cytometry to detect proteins of interest on a single-cell level,^23–25^ and it is one of the suggested models, along with T-cell dependent antibody response (TDAR) to identify compounds of potential immunotoxic risk.^26,27^ Immunotoxicity evaluation is a valuable tool, in combination with other safety endpoints including hematology, clinical chemistry, and histopathology to characterize the toxicity profile of novel food ingredients and was thoroughly addressed in the current work.

Thus this study was designed to assess various parameters of toxicity and immunotoxicity following oral ingestion of effera™. Sprague Dawley rats are a common model used to study immunotoxicity, and humans and rats share immunological similarities making them a suitable model to study immunotoxic effects.^28–31^ Rats are a traditional animal model used to study oral dosing, and the route of dosing is typically aligned with the route of potential human exposure.^31^ The aim of this study was to evaluate safety and toxicity parameters of the test product, effera™, via daily oral administration for 28 days in adult Sprague Dawley rats conducted under good laboratory practice (GLP).^32–34^

## MATERIALS & METHODS

### Regulatory Compliance

All experimental and data collection procedures followed validated methods and standard operating procedures developed by the respective testing facility where methods and procedures were performed: Charles River Laboratories (CRL), Mattawan, MI, USA, Michigan State University (MSU), East Lansing, MI, and Helaina, Inc. (New York, NY, USA). All data was verified by the testing facility or independent scientists at MSU or Helaina, Inc. The study was conducted in accordance with the U.S. Department of Health and Human Services, Food and Drug Administration, United States Code of Federal Regulations, Title 21, Part 58: Good Laboratory Practice for Nonclinical Laboratory Studies and as accepted by Regulatory Authorities throughout the European Union (OECD Principles of Good Laboratory Practice), Japan (MHLW), and other countries that are signatories to the OECD Mutual Acceptance of Data Agreement. Exceptions from the above regulations are as follows: the bLF bioanalytical sample analysis and immunophenotyping analysis were not conducted in accordance with GLP requirements but were conducted in accordance with internal protocols and standard operating procedures. This has no impact on the study results as all results were verified by independent scientists at Helaina Inc, and MSU, respectively.

### Test Substances

The *Komagataella Phaffii* recombinant system was used to produce the active test product, effera™ (effera™, Heliana Inc.), as previously described.^15,22^ Briefly, effera™ was purified by microfiltration/diafiltration and cation exchange chromatography and spray-dried to powder form. Purity was determined to be greater than 98% of the protein content and iron saturation was determined to be 52% saturated. Prior to use, effera™ was reconstituted and diluted into a sodium citrate buffer (pH 5.5). Concentration of the test article was adjusted for purity of rhLF in effera™ powder to ensure proper active concentrations and the quantities tested in this study, 200, 600 and 2000 mg/kg/d, refer to the content of rhLF in effera™.

The active comparative control product, bLF, was isolated from bovine milk and obtained from Lactoferrin Co, Australia (Product 11683). Purity was provided by the supplier and assessed to be greater than 95% and iron saturation to be 9.9% saturated. Prior to use, bLF was reconstituted (adjusted for purity) and diluted into a sodium citrate buffer (pH 5.5) and the active ingredient was equal to 2000 mg bLF/kg/d.

### Animals and Treatment

Animal husbandry and product administration were performed by CRL where 154 adult, 8-week-old Spague Dawley rats (77 male, 77 female) were acclimated for 7 days and then randomly assigned to treatment and control groups using a computer randomization process. Animals were housed in solid-bottom cages (2-3 animals per cage) with nonaromatic bedding and appropriate animal enrichment, adhering to the USDA Animal Welfare Act (9 CFR, Parts 1, 2, and 3), and the Guide for the Care and Use of Laboratory Animals (NRC, Eight Edition).^32^ Animals were fed an ad libitum basal diet of Block Lab Diet® (Certified Rodent Diet #5CR4, PMI Nutrition International, Inc.) and ad libitum treated tap water was provided. During the urinalysis sample collection period, animals were offered ad libitum DietGel® 76A and HydroGel®. Fluorescent lighting was provided 12 hr/day, and temperature (68 – 79°F) and humidity (30 – 70%) of the room was maintained according to the testing facility standard operating procedure (SOP).

The main study for determining the potential toxicity of effera™ test product was conducted in 100 rats where dosing was administered to five groups of 10 rats/sex/dose as follows: sodium citrate buffer vehicle control (Group 1); 200 mg/kg effera™ (Group 2); 600 mg/kg effera™ (Group 3); 2,000 mg/kg effera™ (Group 4); 2,000 mg/kg bLF (Group 5) (**Table 1**). All products were administered through oral gavage once daily for 28 days. Immunotoxicology assessment for T-cell dependent antibody response (TDAR) in the main study animals was also performed by administering 1 mL injections of Keyhole Limpet Hemocyanin (KLH), an antigen for assessing adaptive immune responses, into the animals’ scapular region on Days 2 and 16.

**Table 1.** . Experimental Design (Main Study for Toxicity)

| Group | Test Product | Dose Level<br>(mg/kg/d) | Dose Volume<sup>a</sup><br>(mL/kg) | Dose Concentration<br>(mg/mL) | Toxicity Study |  |
| --- | --- | --- | --- | --- | --- | --- |
|  |  |  |  |  | Males<br>(n) | Females<br>(n) |
| 1 | Vehicle Control | 0 | 10 | 0 | 10 | 10 |
| 2 | effera™ | 200 | 10 | 20 | 10 | 10 |
| 3 | effera™ | 600 | 10 | 60 | 10 | 10 |
| 4 | effera™ | 2000 | 10 | 200 | 10 | 10 |
| 5 | Bovine LF | 2000 | 10 | 200 | 10 | 10 |

<sup>a</sup>Dose volume was based on the most recent body weight measurement.

A secondary satellite study was conducted to determine the toxicokinetic (TK) characteristics of LF and was carried out in an additional 54 rats where dosing was administered to five additional groups as follows: 3 rats/sex/dose received sodium citrate buffer vehicle control (Group 6) and 6 rats/sex/dose received 200 mg/kg effera™ (Group 7); 600 mg/kg effera™ (Group 8); 2,000 mg/kg effera™ (Group 9); 2,000 mg/kg bLF (Group 10) (**Table 2**). All products were administered through oral gavage once daily for 28 days.

**Table 2.** Experimental Design (Secondary Study for Toxicokinetics)

| **Group** | **Test Product** | **Dose Level**<br>**(mg/kg/d)** | **Dose Volume<sup>a</sup>**<br>**(mL/kg)** | **Dose Concentration**<br>**(mg/mL)** | **Toxicokinetic Study<sup>b</sup>** |  |
| --- | --- | --- | --- | --- | --- | --- |
|  |  |  |  |  | **Males**<br>**(n)** | **Females**<br>**(n)** |
| **6** | **Vehicle Control** | **0** | **10** | **0** | **3** | **3** |
| **7** | **effer<sup>TM</sup>** | **200** | **10** | **20** | **6** | **6** |
| **8** | **effer<sup>TM</sup>** | **600** | **10** | **60** | **6** | **6** |
| **9** | **effer<sup>TM</sup>** | **2000** | **10** | **200** | **6** | **6** |
| **10** | **Bovine LF** | **2000** | **10** | **200** | **6** | **6** |

<sup>a</sup>Dose volume was based on the most recent body weight measurement.
<sup>b</sup>Three animals/sex/group were also designated for anti-LF antibody blood sample collection.

### Dose Formulation Analysis

Sample Preparation: Prior to analysis, samples were diluted with 50 mM Tris in water (pH 8) and filtered through a 0.2µm polyethersulfone (PES) filter. Samples underwent a secondary dilution in the same manner to be within the range of the calibration curve. Vehicle samples were also diluted with 50 mM Tris (pH 8) using a dilution factor of 200. An aliquot of each sample was injected into the HPLC-UV system for analysis using the method described below and evaluated at 214 nm.

The conditions used for HPLC-UV analysis included a liquid chromatography system equipped with an Agilent AdvanceBio RP-mAb C4 column, 2.1 x 75 mm, 3.5 μm particle size with a gradient flow of 0.1% trifluoroacetic acid in water (mobile phase A) and 0.09% trifluoroacetic acid in acetonitrile (mobile phase B) at a flow rate of 0.5 mL/minute. Empower™ 2 was used to quantitatively determine the amounts of analytes in samples, including test articles in formulation.

### Animal Observations

All animals were observed cage-side post-dose each day and cage-side observations were made twice daily for morbidity and mortality. Detailed clinical observations and food consumption observations were performed weekly. Body weights were recorded prior to study commencement, weekly, and on the day prior to study termination; ophthalmic parameter observations were performed via indirect ophthalmoscopy by a trained ophthalmologist for deviations from commonly encountered variations. These observations were conducted prior to study commencement and on the day prior to study termination.

### Terminal Procedures

All animals were examined carefully for external abnormalities including palpable masses prior to termination. Animals were euthanized by carbon dioxide inhalation followed by exsanguination or other facility-approved SOP to ensure death. All animals underwent a complete necropsy examination that included evaluation of the carcass and musculoskeletal system. External surfaces and orifices, cranial cavity and external surfaces of the brain, and thoracic, abdominal, and pelvic cavities with their associated organs and tissues were also assessed. The following organs were collected and weighed: brain, epididymis, adrenals, pituitary, prostate, thyroid, heart, kidney, liver, spleen, testis, thymus, ovaries, and uterus/cervix. A veterinary pathologist was available for consultation during scheduled necropsy.

### Hematology

Blood samples were obtained from each main study animal on Day 29 immediately after euthanasia via vena cava venipunctures prior to terminal necropsy. A 1 mL blood sample was collected into K_2_EDTA-coated plastic tubes and underwent hematology measurements. Hematology assessment consisted of leukocyte count (total and absolute differential); erythrocyte count; hemoglobin; hematocrit; absolute reticulocytes; mean corpuscular volume, hemoglobin, hemoglobin concentration (calculated), and red blood cell distribution width; cellular hemoglobin concentration mean; platelet count; mean platelet volume and count; blood smear (preserve and stain); and percent reticulocytes.

### Coagulation

Blood samples were obtained from each main study animal on Day 29 immediately after euthanasia via vena cava venipunctures prior to terminal necropsy. A 1.2 mL blood sample was collected into sodium citrate-coated plastic tubes and processed for plasma coagulation assessment. Coagulation assessment consisted of prothrombin time, activated partial thromboplastin time, fibrinogen, and sample quality.

### Clinical Chemistry

Blood samples were obtained from each main study animal on Day 29 immediately after euthanasia via vena cava venipunctures prior to terminal necropsy. A 1.3 mL blood sample was collected into a serum gel separator tube and serum was subsequently processed for clinical chemistry procedures. Clinical chemistry assessment consisted of alkaline phosphatase, total bilirubin, aspartate aminotransferase, alanine aminotransferase, urea nitrogen, creatinine, total protein, albumin, globulin (calculated), albumin/globulin ratio (calculated), glucose, total cholesterol, triglycerides, electrolytes (sodium, potassium, chloride), calcium, phosphorus, sample quality, unsaturated iron binding capacity (UIBC), total iron binding capacity (TIBC) (calculated), iron, ferritin, and transferrin saturation (TSAT).

### Urinalysis

Urine samples were collected from the main study animals for a period of at least 12 hours, and all available urine samples collected were used to conduct urinalysis measurements. Urinalysis assessment consisted of measurement of volume, color and appearance, specific gravity, pH, protein, glucose, bilirubin, ketones, occult blood, and urobilinogen.

### T-cell Dependent Antibody Response Analysis (TDAR)

Blood samples were obtained from each main study animal in Groups 1 – 5 on Day 1 and Day 15 via the sublingual vein and on Day 29 immediately after euthanasia via vena cava venipunctures prior to terminal necropsy. A 0.5 mL blood sample was collected into plastic tubes and allowed to clot at room temperature for at least 30 min until centrifuged. Serum was then aliquoted into separate tubes and assessed for KLH-induced immunoglobulin G (IgG) and immunoglobulin M (IgM) in accordance with facility SOPs.

### Bioanalytical Assessments

#### Human Lactoferrin ECLIA

An electrochemiluminescent immunoassay (ECLIA) was developed and validated by the testing facility (CRL) to detect human lactoferrin concentration in rat serum ranging from 1.00-200 ng/mL. A minimum required dilution (MRD) of 1:10 was necessary to eliminate matrix effects (e.g. serum). Standards were prepared using effera™ that was used in the oral administration to rats and diluted in rat serum. Positive controls were prepared by spiking known concentrations of effera™ into rat serum. Anti-human lactoferrin antibody-biotin at 0.5 µg/mL in 1X PBS (Novus Biologicals; Cat: LLC NBP2 89794B) working solution was added to streptavidin coated Meso Scale Discovery (MSD®) plates, incubated at ambient temperature with shaking for 30-60 minutes and washed. Prior to sample plating, the plate was blocked for 1-4 hours at ambient temperature with shaking, and then washed. Standards, quality controls, blanks, and required samples were added to the appropriate wells of the plate, and the plate was incubated at ambient temperature with shaking for 1 hour and washed. The goat anti-human lactoferrin antibody (Bethyl Labs; cat A80-143A)-SulfoTag™ (prepared by CRL) detection working solution was added to the wells at 1 µg/mL, and the plate was incubated at ambient temperature with shaking for 1 hour and washed. Finally, 2X MSD® Read Buffer T was added to the wells. Plates were read within 10 minutes of addition of MSD® Read Buffer T. Lactoferrin concentration was interpolated from the standard curve using a 4PL regression model with 1/F^2^ weighting.

#### Bovine Lactoferrin ELISA

A Sandwich ELISA (Bovine Lactoferrin ELISA kit, NBP3-12185, Novus Biologicals) was used and adopted to determine bovine lactoferrin concentrations in rat serum to detect bLF ranging from 0.69 – 500 ng/ml. The assay was performed per the manufacturer’s instructions with the following modifications: standards were prepared using bLF (Lactoferrin Company; Product 11683) that was used in the oral administration to rats, diluted in 1:10 rat serum, positive controls were prepared by spiking known concentrations of bLF into 1:10 rat serum, and the study samples were diluted 1:10 prior to testing. LF concentration was interpolated from the standard curve; limit of quantitation was 6.9-5,000 ng/mL

### Toxicokinetic Parameters

To assess toxicokinetic parameters, a 0.3 mL blood sample was obtained from the sublingual vein at 0.5 and 1.5 h on Day 1 and 28 from satellite animals assigned to the vehicle control group (Group 6) in the secondary satellite study, and at 0.25, 0.5, 1, 1.5, 3, 6, 12, and 24 h for rats assigned to Groups 7 – 10 in the secondary satellite study. Samples were collected into plastic tubes and centrifuged for serum collection within 1 hour of collection. The concentration of LF within serum was determined using the detection methods described above and the results were assessed for toxicokinetics: time to maximum concentration (T_max_), maximum concentration (C_max_), C_max_/dose, area under the curve (AUC)_tlast_/dose, AUC_0-4_, AUC_0-4_/dose, time after last dosing at which the last quantifiable concentration was observed (T_last_), (female: male) F:M AUC, F:M C_max_, AUC from T1 to T2 after repeat dosing divided by the AUC from T1 to T2 during the initial dosing (R_AUC_), and the C_max_ after repeat dosing divided by the C_max_ during the initial dosing interval (R_Cmax_).

### Immunogenicity

#### Anti-Lactoferrin Antibodies in Serum

Two assays were developed and validated at CRL to detect anti-lactoferrin antibodies in rat serum. One assay detected antibodies against bLF, while the other assay detected antibodies against effera™. Following development, assay performance was evaluated for sensitivity, precision, and selectivity. Moreover, screening cut points for each assay were statistically determined to help define the assay sensitivity. The cut point is the level of response above which samples are defined as positive and below which samples are defined as negative for the presence of anti-LF antibodies. During performance evaluation and sample testing, the assays included controls. The assay controls were either pooled rat serum unspiked (negative control; NC) or rat serum spiked with commercially available anti-bLF (Bethyl Labs; Cat A10-126A) or anti-hLF (Bethyl Labs; Cat A80-143A) antibodies, respectively for each assay, at high (high positive control; HPC) and low (low positive control; LPC) levels.

#### Anti-human Lactoferrin Antibody Assay

A bridging ECL assay was used to detect anti-hLF antibodies in rat serum. The assay was designed to first perform a screening assay, followed by a confirmatory assay for any sample above the screening cut point (≥1.04-fold increase above median NC for the plate). For any sample above the cut point of the confirmatory assay (> 32.4% immunodepletion), an additional titer assay was performed with a cut point of ≥1.15-fold increase above median NC. An MRD of 1:25 for each sample was required. For the screening and titer assays, a master mix was prepared with 1.0 µg/mL biotinylated effera™ (biotinylated at CRL) and 1.0 µg/mL effera™-SulfoTag™ (added at CRL). For the confirmatory assay, a confirmatory master mix was prepared with 8.0 µg/mL effera™, 1.0 µg/mL biotinylated effera™, and 1.0 µg/mL effera™-SulfoTag™. For each assay, the appropriate master mix was added to a 96-well storage plate followed by addition of controls (HPC, LPC), and unknown samples and incubated at ambient temperature with shaking for 1 hour. Solution was then transferred to a streptavidin Gold MSD® plate and incubated at ambient temperature with shaking for 1 hour. After incubation, the plate wells were washed, and 2X read buffer (MSD® Read Buffer T) was added to the wells of the plate. The raw signal (ECL units) was measured using an MSD® Sector Imager within 10 minutes of addition of the MSD® Read Buffer T.

#### Anti-bovine Lactoferrin Antibody Assay

A bridging ECL assay was used to detect anti-bLF antibodies in rat serum. The screening cut point was determined to be a 1.01-fold increase above median NC for the plate. Due to this extremely low screening cut point factor, the screening tier was not performed during sample analysis. The assay was designed to start using a confirmatory assay, and for any sample above its cut point (> 13.1% immunodepletion), an additional titer assay with a cut point of ≥1.09-fold increase above NC was performed. An MRD of 1:50 for each sample was required. For the confirmatory assay, a master mix was prepared with 0.868 µg/mL biotinylated bLF from Lactoferrin Co (Product 11683; biotinylated at CRL) and 0.868 µg/mL bLF (Lactoferrin Co; Product 11683)-SulfoTag™ (added at CRL) and 5.0 µg/mL bLF (Lactoferrin Co; Product 11683). For each assay, the appropriate master mix was added to a 96-well storage plate followed by addition of controls (HPC, LPC) and unknown samples and incubated at ambient temperature with shaking for 1 hour. The streptavidin Gold MSD® assay plate was blocked by incubating at ambient temperature with shaking for 1-4 hours, washed and then after the 1-hour incubation, the prepared solution was transferred to the plate and incubated at ambient temperature with shaking for 1 hour. After incubation, the plate wells were washed, and 2X MSD® Read Buffer T was added to the wells of the plate. The raw signal (ECL units) was measured using an MSD® Sector Imager within 10 minutes of addition of the Read Buffer T.

#### Immunophenotyping

After terminal necropsy, spleen samples were collected from the satellite animals on Day 29, blind coded, stored refrigerated (2-8℃) and shipped to the test site at Michigan State University (East Lansing, MI) for exploratory spleen immunophenotyping analysis to determine whether oral administration of effera™ protein to rats produces changes in the frequency of major immune cell types in the spleen as evaluated by flow cytometry. Immunophenotyping was conducted blinded to treatment group and dose level.

#### Tissue processing

Splenocytes were isolated by mechanical disruption of spleen samples and made into single cell suspensions in RPMI 1640 medium supplemented with 5% FCS and penicillin/streptomycin. Red blood cells were lysed using Zap-oglobin-II Lytic Reagent (Beckman Coulter) prior to counting splenocytes on a Z1 Beckman Coulter Counter.

#### Flow Cytometry

A total of 2 x 10^6^ splenocytes were placed in a 96-well round bottom culture plate for surface and intracellular antibody staining. Splenocytes were washed using Hank’s Balanced Salt Solution (HBSS, (pH 7.4)), (Invitrogen) and stained with LIVE/DEAD Fixable Near-IR Dead Cell Stain (Gibco Invitrogen) to assess cell viability. Splenocytes were washed with FACS buffer (1× HBSS (pH 7.5) containing 1% BSA and 0.1% sodium azide) and Cell FcRs were blocked with purified mouse anti-rat CD32 (BD Biosciences). After a 15-minute incubation at 4°C, splenocytes were stained for surface proteins.

To quantify Granulocyte/Macrophage/Dendritic cell and B cell populations, the following antibodies were used for immunophenotyping detection: CD45 (OX-1) (Biolegend), CD172a (clone OX-41) (BD Bioscience), CD11b (clone WT.5) (Biolegend), CD25 (clone OX-39) (Biolegend), and AffiniPure Anti-Rat IgG + IgM (Jackson ImmunoResearch). To quantify the T cell populations, the following antibodies were used: CD45 (OX-1) (Biolegend), CD3 (clone 1F4) (Biolegend), CD4 (clone W3/25) (Biolegend), CD8 (clone OX-8) (Biolegend) and CD25 (clone OX-39) (Biolegend).

For FoxP3^+^ T cells: Splenocytes were incubated at 4°C and washed three times with FACS buffer and fixed with FoxP3/Transcription Factor Staining Buffer Set (Invitrogen) fixative and stored at 4°C until intracellular staining. To measure intracellular FoxP3, splenocytes were washed and incubated with 1x Permeabilization buffer solution (Invitrogen) for 15 minutes, then incubated with anti-rat FoxP3 (Invitrogen) for 30 minutes. Splenocytes were washed 3X with the permeabilization buffer and then washed one more time with FACS buffer and resuspended in FACS buffer. Flow cytometric analysis was performed on a Cytek Northern Lights full spectral analyzer (Cytek Biosciences) and analyzed using FlowJo v10.9.0 (Tree Star, Ashland, OR) software.

### Statistical analysis

Statistical analyses were conducted with Provantis version 9. Raw data was tabulated within each time interval, and the mean and standard deviation (or incidence counts for categorical variables) were calculated for each endpoint by sex and group. For each endpoint, treatment groups were compared to the control group. The groups were compared using an overall one-way ANOVA F-test if Levene’s test was not significant, or the Kruskal-Wallis test if it was significant. If the overall F-test or Kruskal-Wallis test was found to be significant, then pairwise comparisons were conducted using Dunnett’s or Dunn’s test, respectively. Results of all pairwise comparisons were reported at the 0.05 and 0.01 significance levels. All endpoints were analyzed using two-tailed tests unless indicated otherwise.

## RESULTS

### Dose Formulation Analysis

The results of the dose formulation analysis of the study samples are presented in **Table 3**. All dose formulation results were within ± 10%, for effera™, and ± 15%, for bLF, of the target concentration, and within ≤ 5% relative standard deviation (RSD). Vehicle control samples were below the limit of quantitation. These analytical data demonstrated acceptable performance of the methods for all reported results, and that dose formulations were prepared at the intended target LF concentration and were homogeneous.

**Table 3.** Formulation Analysis of effera™ and Bovine LF

| **Test Product** | **Dose Level**<br>**(mg/kg/day)** | **Nominal**<br>**Concentration**<br>**(mg/mL)** | **Average Calculated**<br>**Concentration**<br>**(mg/mL)\*** | **Average %**<br>**Recovery<sup>a</sup>** | **%Relative**<br>**Standard**<br>**Deviation** |
| --- | --- | --- | --- | --- | --- |
| **Vehicle Control** | **0** | **0.000** | **BLQ** | **NA** | **NA** |
| **effer<sup>TM</sup>** | **200<sup>b</sup>** | **20.000** | **19.007 – 20.077** | **95.0 – 100.4** | **1.965 – 2.206** |
| **effer<sup>TM</sup>** | **600** | **60.000** | **59.300 – 59.475** | **98.8 – 99.1** | **0.075 – 0.970** |
| **effer<sup>TM</sup>** | **2000<sup>b</sup>** | **200.000** | **195.749 – 200.252** | **97.9 – 100.1** | **0.637 – 1.482** |
| **Bovine LF** | **2000<sup>b</sup>** | **200.000** | **198.121 – 212.644** | **99.1 – 106.3** | **1.546 – 4.747** |

**\*Concentration refers to active LF (rhLF or bLF) in formulation**
**Abbreviations: BLQ, below the limit of quantitation (<4.0000 mg/mL); LF, lactoferrin**
**<sup>a</sup>Average % recovery was calculated from the nominal concentration.**
**<sup>b</sup>The averaged result from homogeneity analysis served as concentration verification.**

### Clinical Observations

All animals survived to scheduled termination, and no effects on clinical findings were observed in the effera™ or bLF-treated groups (**Supplementary Table 1**). There were no effera™ or bLF-related effects on terminal ophthalmology examinations (**Supplementary Table 1**).

### Body Weights and Food Consumption

The mean body weights for effera™ and bLF-treated groups were comparable to the vehicle control group throughout the study and were unaffected by treatment (**Figure 1**). Mean body weight gains were also not significantly different between the vehicle control group and test product groups for both males and females (**Table 4**). Mean food consumption intake was also not significantly affected by treatment. However, there was a non-significant trend for intake to be lower at 2000 mg effera™/kg/d for most measurement intervals for both males (–4.4% to -8.8%) and females (−4.6% to -6.3%) when compared to the vehicle control group, with the exception of the Day 8 – 15 intervals for females which was higher than the vehicle control group (+23.0%) (**Supplementary Table 1**). Similarly, mean food consumption at 2000 mg bLF/kg/d was lower for all measurement intervals for both males (−4.5% to -8.2%) and females (−3.8% to –11.3%) when compared to the vehicle control group.

**Figure 1.**
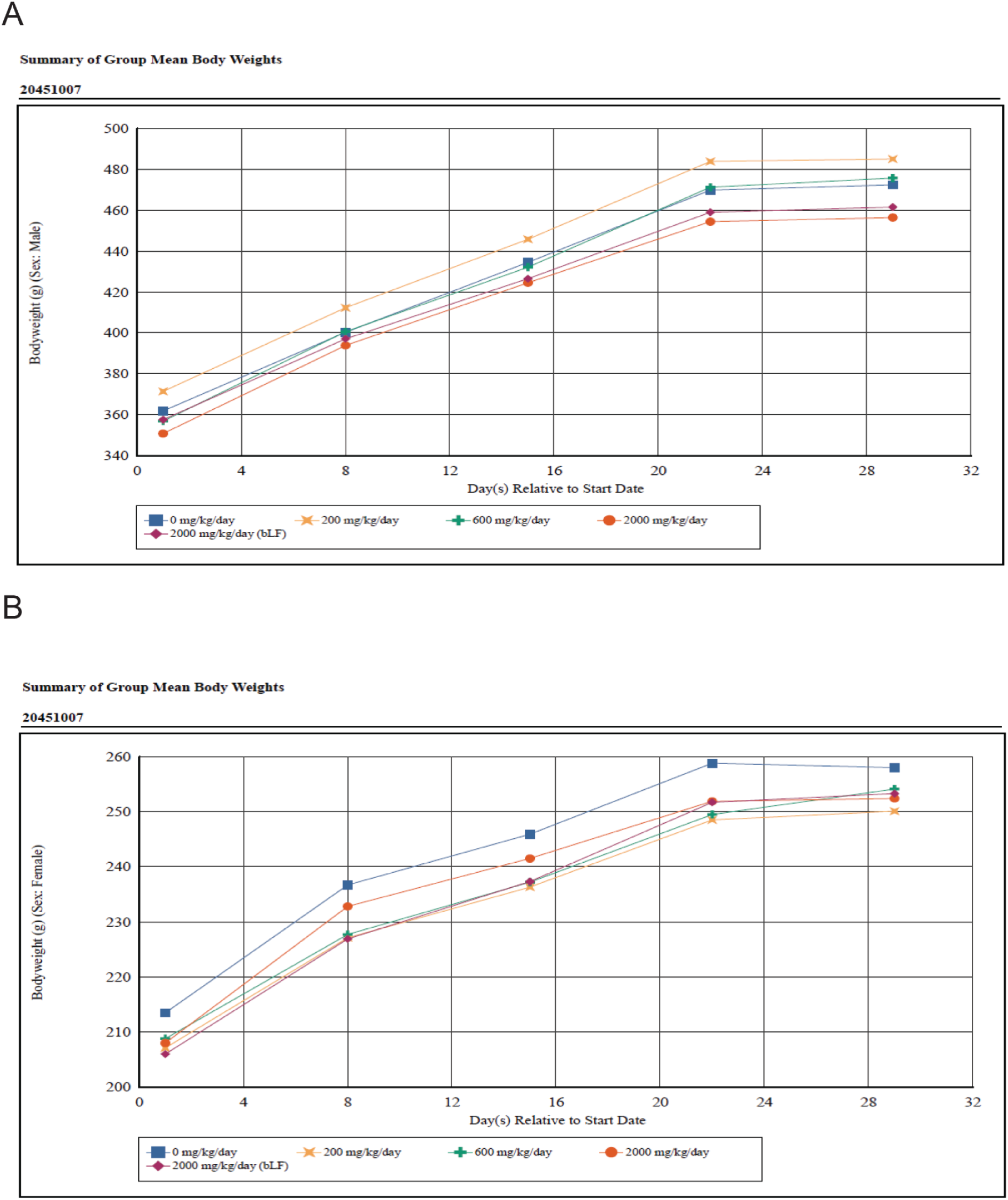
Body weight

**Table 4.** Body Weight Gain from Day 1 - 29

| **Group** | **Test Product** | **Dose Level**<br>**(mg/kg/d)** | **Mean Weight Gain (SD)** |  |
| --- | --- | --- | --- | --- |
|  |  |  | **Males** | **Females** |
| **1** | **Vehicle Control** | **0** | **110.4 (20.8)** | **44.5 (7.1)** |
| **2** | **effera™** | **200** | **113.6 (12.7)** | **43.0 (9.1)** |
| **3** | **effera™** | **600** | **118.8 (29.0)** | **45.3 (11.4)** |
| **4** | **effera™** | **2000** | **105.6 (14.9)** | **44.4 (7.9)** |
| **5** | **Bovine LF** | **2000** | **103.9 (13.5)** | **47.3 (11.3)** |

**Abbreviations: SD, standard deviation; n = 10 for animals/sex/group**

### Hematology and Coagulation

There were no significant effera™ or bLF-related effects on measured hematology parameters in either sex at any dose level at termination (**Supplementary Table 1**). There were no effera™ or bLF-related effects on measured coagulation parameters in either sex at any dose level at termination (**Supplementary Table 1**).

### Clinical Chemistry

There were no significant effera™ or bLF-related effects on measured clinical chemistry parameters in either sex at any dose level at termination (**Supplementary Table 1**). On Day 29, prior to termination, there was a non-significant tendency for a 29% higher cholesterol concentration, 8% higher globulin concentration and 4% higher total protein concentration and a 7% lower albumin to globulin ratio in males treated with 2000 mg effera™/kg/d compared to the vehicle control. In males treated with 2000 mg bLF/kg/d, there was a non-significant tendency for 27% lower triglyceride concentration. On Day 29, in females at 2000 mg effera™/kg/d there was a non-significant 33% higher mean serum ferritin concentration.

### Urinalysis

There were no effera™ or bLF-related effects on measured urinalysis parameters in either sex at any dose level at termination with the exception of urine specific gravity (USG). On Day 29, mean urine specific gravity values for males and females at 2000 mg effera™/kg/d were 59% (p ≤ 0.01) and 24% (p ≤ 0.05) higher than the control for males and females, respectively.

### T-Cell Dependent Antibody Response Analysis

Administration of effera™ appeared to decrease IgG concentration in males at 200 mg/kg/d on Day 15 by 44% and Day 29 by 33% compared to vehicle control-treated males (**Table 5**). Effera™ treatment increased IgG concentration in females at 2000 mg/kg/d on Day 15 by 111% compared to vehicle control-treated females. This was attenuated to a more modest 57% increase compared to vehicle control in IgG concentration by Day 29. Similarly, administration of bLF did not alter the Anti-KLH IgG response in males at 2000 mg/kg/day on Day 15 and Day 29 and showed increased Anti-KLH IgG concentration in females at 2000 mg/kg/day on Day 15 and Day 29 (85% and 78%, respectively) compared to the vehicle control group females; however, none of these results were statistically significant (**Supplementary Table 1)**.

**Table 5.** T-Cell Dependent Antibody Response Changes by Test Product

|  |  |  | IgG |  |  |  |  |  | IgM |  |  |  |  |  |
| --- | --- | --- | --- | --- | --- | --- | --- | --- | --- | --- | --- | --- | --- | --- |
|  |  |  | Male |  |  | Female |  |  | Male |  |  | Female |  |  |
| Group | Test Product | Dose Level (mg/kg/d) | Pre-Dose | Day 15 | Day 29 | Pre-Dose | Day 15 | Day 29 | Pre-Dose | Day 15 | Day 29 | Pre-Dose | Day 15 | Day 29 |
| 2 | effera™ | 200 | — | 0.56 | 0.67 | — | — | — | 0.37 | 0.48 | 0.82 | — | — | — |
| 3 | effera™ | 600 | — | — | — | — | — | — | 0.47 | 0.77 | 1.43 | — | — | — |
| 4 | effera™ | 2000 | — | — | — | — | 2.11 | 1.57 | 1.74 | 0.70 | 1.16 | 2.94 | 1.11 | 0.74 |
| 5 | bLF | 2000 | — | — | — | — | 1.85 | 1.78 | 1.43 | 0.50 | 0.78 | 2.08 | 1.44 | 0.65 |

Abbreviations: bLF, Bovine Lactoferrin; IgG, Immunoglobulin G; IgM, Immunoglobulin M; n = 10 for animals/sex/group.
A dash (—) indicates absence of a test article-related change. Numerical values indicate fold change of the treated group mean value relative to the control group, pre-treatment mean value. Bold indicates statistical differences from 0 mg/kg/day, p < 0.01.

Neither effera™ nor bLF administration significantly affected IgM concentration in either sex at Day 15 or Day 29 compared to sex-matched vehicle controls (**Supplementary Table 1)**. However, within females assigned to either the 2000 mg effera™/kg/d or 2000 mg bLF/kg/d group, there was a mild but statistically significant higher pre-dose IgM concentration of 194% and 108%, respectively, compared to females assigned to the vehicle control group.

### Immunophenotyping results

Oral gavage with LF did not alter spleen weight, and there was no difference in spleen cellularity observed in any treatment group. Oral gavage with effera™ or with bLF did not have any significant effect on cell viability in either female or male rats in any of the treatment groups. Additionally, no treatment-related effects were observed in the overall composition of the immune cell repertoire or at the population level, including CD172^+^ myeloid-derived immune cells, granulocytes, monocytes, macrophages, dendritic cells, B cells, T cells (CD4^+^, CD8^+^, regulatory CD4^+^), NK, and NK T cells (**Table 6**). Likewise, while there were gross differences in the numbers of specific immune cell populations between male and female rats, there were no sex-specific differences related to oral administration of either effera™ or bLF.

**Table 6.** Immunophenotyping Results in Male and Female Rat Spleen Following Oral Gavage Administration of effera™ or bLF

|  | Female |  |  |  |  | Male |  |  |  |  |
| --- | --- | --- | --- | --- | --- | --- | --- | --- | --- | --- |
|  | 0 | 200<br>effera™ | 600<br>effera™ | 2000<br>effera™ | 2000<br>bLF | 0 | 200<br>effera™ | 600<br>effera™ | 2000<br>effera™ | 2000<br>bLF |
| Total Splenocytes<br>(x10<sup>6</sup> per spleen) | 7.57<br>(0.78) | 5.45<br>(1.01)* | 6.77<br>(0.92) | 5.32<br>(0.88)* | 6.78<br>(1.35) | 14.67<br>(4.08) | 9.12<br>(2.42) | 11.97<br>(4.57) | 10.87<br>(2.11) | 10.35<br>(2.84) |
| Gran/Mono/<br>Mac/DC | 13.20<br>(4.23) | 13.45<br>(2.05) | 14.85<br>(3.92) | 13.08<br>(2.42) | 12.02<br>(1.03) | 8.37<br>(1.98) | 9.07<br>(1.35) | 10.25<br>(3.09) | 9.88<br>(1.52) | 10.10<br>(1.3) |
| Macrophages<br>(Within G/M/M/DC) | 28.43<br>(6.97) | 31.28<br>(4.31) | 33.48<br>(5.81) | 31.34<br>(5.21) | 34.72<br>(3.78) | 38.78<br>(8.44) | 2928<br>(5.66) | 34.17<br>(9.16) | 32.45<br>(6.58) | 34.68<br>(7.86) |
| Dendritic Cells<br>(within Mac) | 5.57<br>(1.11) | 6.07<br>(1.92) | 5.73<br>(1.09) | 5.56<br>(1.65) | 5.57<br>(1.34) | 2.93<br>(1.17) | 3.27<br>(1.36) | 3.38<br>(0.86) | 3.43<br>(1.05) | 3.67<br>(1.38) |
| B Cells | 60.00<br>(2.65) | 58.82<br>(2.16) | 57.02<br>(3.14) | 58.88<br>(3.36) | 61.00<br>(1.87) | 57.87<br>(1.82) | 59.68<br>(4.97) | 59.8<br>(2.22) | 60.42<br>(4.22) | 59.90<br>(3.73) |
| T Cells | 26.07<br>(5.21) | 24.17<br>(3.16) | 24.13<br>(1.78) | 25.96<br>(1.85) | 23.15<br>(1.69) | 30.27<br>(8.34) | 26.55<br>(3.91) | 25.43<br>(3.14) | 25.80<br>(3.12) | 27.42<br>(5.56) |
| CD4<sup>+</sup> T Cells<br>(Within T Cells) | 54.63<br>(0.76) | 56.88<br>(5.03) | 51.23<br>(1.87) | 52.38<br>(5.17) | 54.22<br>(3.29) | 55.67<br>(2.90) | 49.87<br>(3.90) | 55.85<br>(6.65) | 53.17<br>(5.43) | 54.88<br>(5.07) |
| Treg Cells<br>(Within CD4<sup>+</sup> T cells) | 9.00<br>(0.95) | 9.97<br>(1.24) | 10.12<br>(0.85) | 10.58<br>(1.16) | 11.10<br>(1.40) | 8.73<br>(2.02) | 10.05<br>(0.49) | 9.03<br>(0.86) | 9.02<br>(0.95) | 10.27<br>(1.05) |
| CD8<sup>+</sup> T Cells<br>(Within T Cells) | 36.47<br>(3.89) | 36.47<br>(6.29) | 36.27<br>(2.90) | 37.72<br>(4.70) | 34.77<br>(4.90) | 36.83<br>(2.63) | 34.32<br>(3.71) | 31.28<br>(5.47) | 34.93<br>(2.62) | 33.62<br>(4.68) |
| Activated T Cells<br>(Within T Cells) | 20.00<br>(4.28) | 21.98<br>(2.86) | 22.48<br>(2.09) | 20.90<br>(2.22) | 22.43<br>(3.28) | 16.03<br>(2.48) | 17.90<br>(2.65) | 21.33<br>(6.37) | 19.07<br>(3.03) | 19.03<br>(1.90) |
| Natural Killer Cells | 12.13<br>(1.55) | 13.77<br>(2.19) | 13.30<br>(1.82) | 12.98<br>(2.13) | 12.27<br>(1.27) | 12.00<br>(0.82) | 10.98<br>(1.90) | 11.8<br>(1.82) | 11.27<br>(1.23) | 11.03<br>(1.08) |
| NKT Cells<br>(Within T Cells) | 10.53<br>(3.25) | 12.67<br>(2.65) | 12.07<br>(1.70) | 10.56<br>(1.85) | 10.75<br>(1.65) | 16.60<br>(3.93) | 17.23<br>(6.09) | 17.95<br>(4.72) | 16.47<br>(2.32) | 16.92<br>(1.79) |

Doses are in mg/kg/day; N = 3 vehicle (0) group (sodium citrate buffer), N = 6 treatment groups. Female group 2000 mg/kg/day effera™ N = 5. Spleen cell populations are presented as percent ± (SD).
\* = Significance at p < 0.05.
bLF = bovine lactoferrin, DC = dendritic cells; G = granulocytes, M = monocytes/macrophages

### Immunogenicity results

Regarding anti-effera™ antibody evaluation, control group animals were generally negative for the presence of anti-effera™ antibodies on Days 1 and 29 (**Supplementary Table 2**). The exception was for one female animal for which anti-effera™ antibodies were detected on Days 1 and 29.

Anti-effera™ antibodies were detected in 1 out of 6 (17%) and 4 out of 6 (67%) animals from the 200 mg/kg on Days 1 and Day 29, respectively. Among the 600 mg/kg treatment group, anti-effera™ antibodies were detected in 4 out of 6 (67%) animals on Day 29, and 6 out of 6 (100%) animals had anti-effera™ antibodies in the 2000 mg/kg by Day 29. In the comparative group, anti-bLF antibodies were detected in one out of 6 (17%) animals on Day 29 (**Supplementary Table 3**).

### Toxicokinetic Evaluation

Results indicated that effera™ was absorbed rapidly and could be detected in serum within 15 minutes of administration for males dosed at all levels on both Day 1 and Day 28 (**Supplementary Figure 1A**). Levels of bLF were detected in males up to 1.5 hours post-dose on Day 1, with the highest concentration observed at 30 minutes post-dose and quickly cleared from serum (**Supplementary Figure 1A**). On Day 28, bLF was only detected at the 15-minute time point in males (**Supplementary Figure 1B**), indicating rapid absorption and clearance. On Day 1, detection of effera™ in female rats was observed within 15 minutes (**Supplementary Figure 1C**). However, greater variability was observed across doses compared to male rats with levels peaking for low dose at 1.5 hours and the mid and high doses at 30 minutes. On Day 28 for the low dose females, effera™ could be detected by 15 minutes and was quickly cleared, low levels of effera™ were detected for the mid dose females; whereas much higher levels of effera™ were detected in the high dose group (**Supplementary Figure 1D**). Furthermore, these results show higher overall detectable levels of effera™ in the female rats. Detection of bLF was limited to 15-30 minutes in the female rats on both Day 1 and Day 28, with higher levels at 15 minutes and increased levels on Day 28 compared to Day 1 (**Supplementary Figure 1C, D**).

The toxicokinetic evaluation yielded limited concentration results, and high variability existed among the detected values. Furthermore, systemic exposure to effera™ appeared to be sex dependent such that individual effera™ serum concentrations, C_max_, and AUC values were greater in females than in males (F:M C_max_ ratios ranged from 0.108 to 56.2 and F:M AUC_0-24hr_ ratios ranged from 0.485 to 13.7); therefore, data for females and males were reported separately. Specifically, four male animals at the 200 mg/kg had serum concentrations below the bioanalytical limit of quantitation (<1.00 ng/mL) at all timepoints on either Day 1 and/or Day 28, and one male animal at the 600 mg/kg on Day 28 was below detection limits.

Following effera™ administration via oral gavage once daily to male rats on Day 1, C_max_ remained similar from 200 to 600 mg/kg and increased from 600 to 2000 mg/kg (**Table 7**). AUC_0-24hr_ values of effera™ did not appear to change from 600 to 2000 mg/kg. A 1:3-fold increase in effera™ dose from 200 to 600 mg/kg resulted in an approximate 1:3.2-fold increase in effera™ C_max_ values. On Day 28, C_max_ increased from 200 to 600 mg/kg and slightly decreased from 600 to 2000 mg/kg whereas AUC_0-24hr_ values of effera™ increased from 600 to 2000 mg/kg. A 1:3:10-fold increase in effera™ dose resulted in an approximate 1:61.0:54.0-fold increase in effera™ C_max_ values and an approximate 2.1-fold increase in effera™ AUC_0-24hr_ values from 600 to 2000 mg/kg.

**Table 7.** Toxicokinetic Parameters of effera™ in Male and Female Rat Serum Following Oral Gavage Administration of effera™

| Day | Sex | Dose (mg/kg) | C<sub>max</sub> (ng/mL) | t<sub>max</sub> | t<sub>last</sub> | AUC<sub>0-24h</sub> | F:M AUC<sub>0-24h</sub> | F:M C<sub>max</sub> | RAUC | RC<sub>max</sub> |
| --- | --- | --- | --- | --- | --- | --- | --- | --- | --- | --- |
| 1 | F | 200 | 851 | 1.5 | 24 | 1270 | 13.7 | 56.2 | NA | NA |
|  |  | 600 | 71.2 | 0.5 | 24 | 61.3 | 0.485 | 3.79 | NA | NA |
|  |  | 2000 | 362 | 0.5 | 24 | 214 | 1.75 | 6.13 | NA | NA |
| 1 | M | 200 | 15.2 | 24 | 24 | NA* | NA* | NA | NA | NA |
|  |  | 600 | 18.8 | 24 | 24 | 126 | NA | NA | NA | NA |
|  |  | 2000 | 59.1 | 0.25 | 24 | 122 | NA | NA | NA | NA |
| 28 | F | 200 | 39.5 | 0.25 | 3 | NA* | NA* | 28.2 | NA* | 0.0464 |
|  |  | 600 | 9.22 | 1.5 | 24 | NA* | NA* | 0.108 | NA* | 0.130 |
|  |  | 2000 | 1530 | 0.25 | 24 | 511 | 6.08 | 20.2 | 2.39 | 4.23 |
| 28 | M | 200 | 1.40 | 0.5 | 3 | NA* | NA | NA | NA* | 0.0924 |
|  |  | 600 | 85.4 | 0.5 | 3 | 39.5 | NA | NA | 0.313 | 4.55 |
|  |  | 2000 | 75.7 | 0.5 | 24 | 84.1 | NA | NA | 0.689 | 1.28 |

**Abbreviations:** F, Female; F:M = AUC<sub>0-24hr</sub> Female/AUC<sub>0-24hr</sub> Male; F:M = C<sub>max</sub> Female/C<sub>max</sub> Male; M, Male; RAUC = AUC<sub>0-24hr</sub> Day 28/AUC<sub>0-24hr</sub> Day 1; RC<sub>max</sub>, C<sub>max</sub> Day 28/C<sub>max</sub> Day 1
\*AUC is not reported due to less than 3 consecutive quantifiable concentrations.

At 200 mg/kg, C_max_ value decreased in male rats following repeated administration of effera™ on Day 28 compared to Day 1 with a ratio of 0.0924. At 600 mg/kg, C_max_ value increased in male rats on Day 28 compared to Day 1 with a ratio of 4.55 and AUC_0-24hr_ value decreased following repeated administration of effera™ with a ratio of 0.313. At 2000 mg/kg, systemic exposure (C_max_ and AUC_0-24hr_ values) to effera™ in male rats did not appear to change following repeated administration of effera™ with C_max_ and AUC_0-24hr_ accumulation ratios of 1.28 and 0.689, respectively.

Following effera™ administration via oral gavage once daily to female rats on Day 1, C_max_ and AUC_0-24hr_ values decreased from 200 to 600 mg/kg and increased from 600 to 2000 mg/kg (**Table 7**). A 1:3.3-fold increase in effera™ dose from 600 to 2000 mg/kg resulted in an approximate 1:5.1-fold increase in effera™ C_max_ values and an approximate 1:3.5-fold increase in effera™ AUC_0-24hr_ values. On Day 28, C_max_ decreased from 200 to 600 mg/kg and increased from 600 to 2000 mg/kg. A 1:3.3-fold increase in effera™ dose from 600 to 2000 mg/kg resulted in an approximate 1:166-fold increase in effera™ C_max_ values.

At 200 and 600 mg/kg, C_max_ values decreased in female rats following repeated administration of effera™ on Day 28 compared to Day 1 with a ratio of 0.0464 and 0.130, respectively. At 2000 mg/kg, systemic exposure (C_max_ and AUC_0-24hr_ values) to effera™ in female rats increased following repeated administration of effera™ with C_max_ and AUC_0-24hr_ accumulation ratios of 4.23 and 2.39, respectively.

Anti-effera™ antibodies were not determined in all animals and due to the high variability observed in results on Days 1 and 29, no correlation could be obtained in systemic exposure (C_max_ and AUC_0-24hr_ values) to effera™ observed in anti-effera™ antibody positive animals. Limited available data on sex differences in systemic exposure or changes in systemic exposure to bLF following repeated administration of bLF determined that no correlation could be determined in systemic exposure (Cmax and AUC0-24hr values) to bLF observed in the anti-bLF antibody positive animal.

### Organ Weights

There were no significant effera™ or bLF-related effects on measured organ weights for either sex (**Supplementary Table 1**).

## DISCUSSION/CONCLUSION

This 4-week GLP study represents the second study to assess the effect of a novel human lactoferrin produced by *K. phafii* developed by Helaina Inc., on various safety parameters including immunotoxicity. There were no overt differences following 28 days of oral ingestion of up to 2000 mg effera™/kg/d, compared to a citrate buffer negative control and bovine LF comparative control, on various toxicology outcomes, including immunotoxicology parameters in adult Sprague Dawley rats. The test dose of 2000 mg effera™/kg/d is effectively equivalent to ∼400 times the estimated daily intake of rhLF at the 90th percentile of a commercially available product for adult humans.

There were no animal mortalities throughout the entire study period, and the clinical and ophthalmology findings observed in the treated groups were either similar to those observed in the vehicle control group or were infrequent and considered common in Sprague Dawley rats and determined to be unrelated to effera™ or bLF. Mean body weight and body weight gain in both male and female rats were also within normal range for the animal age/sex group, further demonstrating that both the effera™ test product and the bLF comparative control product did not overtly affect animal anthropometrics. Although measured food intake was lower in both the 2000 mg effera™/kg/d and 2000 mg bLF/kg/d treated animals in both males and females. These differences in the food consumption were not statistically significant, but due to the presence in both males and females of the high dose groups, they may be related to increased overall protein intake rather than LF treatment, whether bovine or human. Due to the low magnitude of the differences from the control group and the absence of corresponding significant effects on mean body weights, the slightly lower food consumption at 2000 mg/kg/day is likely not adverse.

All groups, regardless of treatment assignment, had similar hematology, coagulation and clinical chemistry outcomes indicating that effera™ does not adversely affect these parameters. A preceding pilot study was conducted by Peterson et al.^22^ The study concluded that minimal to moderate changes in hematology and chemistry outcomes were observed, including those related to iron homeostasis.^22^ LF is known as an iron-binding protein, and its immunomodulatory effects have been attributed, in part, to its iron-binding or iron-chelating properties.^18,35,36^ For example, LF supplementation has been shown to improve iron status and decrease respiratory tract infection in infants.^37,38^ In addition, LF has been reported to exert an anti-inflammatory effect against interleukin 6 which is also known to support iron homeostasis.^39^ This present study did not find any significant effects on any iron-related parameters assessed. Of note, the products used in both studies were similar in iron saturation (>50% for rhLF and ∼10% for bLF) and therefore, iron status was unlikely to have played a role in these differences. However, these observations may be explained in a number of other ways. For example, due to LF having a “normalizer” effect on iron status when inflammatory anemia is present,^40^ versus simply an additive effect on iron levels. Alternatively, the absorption of non-heme iron has been found to be lower in high iron-bioavailable diets and higher in low iron-bioavailable diets,^41^ subsequently, a diet-based adaptation may also be affecting iron status and furthermore, these are healthy animals with no underlying inflammatory and/or iron-related issues. Thus, an effect on iron homeostasis may not always manifest after LF supplementation.

Our preliminary study also concluded that a similar toxicological profile between effera™ and bLF could be observed, but that these changes did not appear to affect the overall health of the animals.^22^ Bovine LF is commonly used in infant formulas and is commercially available in various supplements.^42^ Consistently, this present study did not find any significant differences between the treatment groups for any clinical chemistry parameter. The present study was conducted over a longer time period (28 d versus 14 d), with a greater number of animals in each treatment group, allowing for greater statistical power. Of note, there were non-significant trends for higher cholesterol, globulin, and total protein concentration in males treated with 2000 mg effera™/kg/d and a higher serum ferritin concentration for females receiving this same treatment and dose. LF supplementation in various animal models and humans have been shown to affect cholesterol metabolism and plasma protein levels, although the exact mechanisms and implications are an active area of study.^43,44^

Urinalysis-based toxicology assessments also found no significant adverse effect of LF treatment, regardless of source or dose, compared to the vehicle control. The mean urine specific gravity values for both sexes treated with the 2000 mg effera™/kg/d were slightly higher than the respective vehicle controls, and its relation to effera™ treatment cannot be ruled out. Urine specific gravity assesses hydration status,^45^ and the inherently high protein (LF) intake of 2000 mg/kg/d may have a dehydration effect. However, the small and inconsistent magnitude of difference and effect among cohort group members, absence of correlative findings in related clinical pathology parameters, and high variability inherent to urinalysis parameters from pan-collected urine specimens preclude definitive interpretation.

TDAR assays are functional assays used in immunopharmacology and immunotoxicity studies to assess the antibody response of IgM and IgG to antigen sensitization (e.g., KLH), and is a standard method to determine the effect of a novel component on the immune system.^46^ In this study, the TDAR assay results indicate neither effera™ nor bLF induce IgG- and IgM-mediated immunotoxicity, and all doses assessed were comparable to vehicle control with no reduction in antibody response against KLH that would be indicative of immunotoxicity. Of note, within females assigned to either the 2000 mg effera™/kg/d or 2000 mg bLF/kg/d group, there was a mildly higher pre-dose IgM concentration versus the vehicle control group; however, these results are pre-dose values and therefore, are not LF-dependent, and may indicate an outcome related to the housing facility and/or animal husbandry.

As an additional measure of immunotoxicity, evaluation of immune cell populations in the spleen using immunophenotyping showed no treatment-related changes in spleen cellularity or spleen-resident immune cell populations in male or female rats in any treatment group. This would suggest that oral administration of effera™ does not affect the number or activation status of the specific cell populations evaluated in the study. The previous study conducted on effera™ showed minor alterations in immune cell populations following 14-day exposure that were not fully recapitulated in this longer-term study. This may be due to adaptability during the 28-day period as described above regarding iron in the diet^19^ and the known effects of iron on the immune system. In the previous study, an increase in activated CD4+ T cells was observed following the administration of both rhLF and bLF suggesting a specific response against LF. In the current study, while no changes were observed in the activated T cells at the 28-day timepoint, the development of anti-rhLF antibodies indicates that there was immune activation specific to rhLF administration.

Lactoferrin absorption results indicate that both effera™ and bLF were absorbed quickly in both male and female rats, with slightly higher levels observed in female rats. Furthermore, while there are observed differences in the absorption between effera™ and bLF, it is important to note that detection was performed using different methods that may have accounted for this variability and due to limitations in positive results, direct comparisons between effera™ and bLF could not be performed. Regarding TK parameters, systemic exposure to effera™ appeared to be sex dependent and was generally greater in females.

However, due to lack of detectability in a number of samples and the high degree of variance when effera™ was detected, results should be interpreted with caution. The C_max_ and AUC_0-24hr_ values resulting from the daily exposure of both males and females to 200 mg/kg and 600 mg/kg effera™ appeared to be generally lower on Day 28 compared to Day 1 whereas at 2000 mg/kg systemic exposure remained approximately the same in males and was higher in females on Day 28 compared to Day 1. This suggests that there is an adaptive toxicokinetic response to the effera™ such that over a period of time, the test product is metabolized more quickly than at initial exposure. However, this response is not present in the highest 2000 mg/kg/day dose indicating that this dose is higher than the toxicokinetic adaptability phase although further research is needed, particularly since limited data did not allow for comparison to the bLF comparative control.

Regarding immunogenicity, anti-effera™ antibodies could not be determined in all animals; however, there was a trend towards an increase related to dose. Of note, there was a positive response for rhLF in one female control animal on both Day 1 and 29. Anti-rhLF antibodies were detected in the same animal on both days and numeric values were three times higher at Day 29 versus Day 1. Although all laboratory protocols were strictly conducted, we cannot rule out exposure to LF prior to and/or during the study period. However, since there was a high degree of variability observed in results on Days 1 and 28 and no correlation to systemic exposure could be determined, further studies may be warranted. Only one bLF animal had a positive anti-bLF antibody response; however since systemic exposure of bLF could not be assessed due to insufficient quantifiable concentrations on both occasions, correlation could not be determined in systemic exposure (Cmax and AUC0-24hr values) to bLF observed in this anti-bLF antibody positive animal. The higher number of antibody positive animals in the 2000 mg/kg effera™ group compared to the 2000 mg/kg bLF group could be due to the relative increased absorption of effera™ and does not necessarily indicate greater immunogenicity would occur in a clinical setting. It is important to note that these results were only used to better understand the TK results of this study. Furthermore, these data support that true immunogenicity potential of a protein food ingredient following ingestion needs to be evaluated in the clinical setting as previously reported.^47^ To this end, a recent clinical study conducted by the authors concluded that effera™ was non-immunogenic (i.e. non-alloimmunizing) in the clinic and safe for use.^48^

Many previous toxicology studies have shown rhLF to be safe and well-tolerated in the rat model, and this study further confirms these results. In addition, this is the first study of its kind to thoroughly evaluate the immunotoxicity potential of any rhLF. These data along with the previous toxicology studies indicate that rhLF is a promising candidate for a safe food ingredient.

In conclusion, in all treatment groups, regardless of dose assignment, no significant adverse effects were observed for body weight, food intake, clinical observations, hematology, coagulation, clinical chemistry, urinalysis, toxicokinetics and immunotoxicological (i.e. immunophenotyping and TDAR) outcomes in male and female Sprague Dawley rats treated with effera™ or bLF for 28 days compared to vehicle control. This study demonstrates that effera™ intake of up to 2000 mg rhLF/kg/d for 28 days is comparable in all assessed safety and toxicology measures to the same dose of bLF and vehicle control and is safe and tolerable with a no observed adverse effect level (NOAEL) of 2000 mg rhLF/kg/d, the highest dose evaluated.

## ACKNOWLEDGEMENTS

The authors would like to thank technical writer Orsolya M. Palacios, RD, PhD for assistance with the preparation of the manuscript, June S. Sass, BS, Yanisa Anaya, BE and Raysa Rosario Martinez, BS for their technical expertise in bovine lactoferrin ELISA methodology.

## FOOTNOTES

Funding: Privately funded by Helaina, Inc.

Declaration of Conflicting Interest: Authors RP, AJC, and C-AM are all employees of Helaina, Inc. Author NEK is a paid consultant of Helaina, Inc.

### Data Availability

Not applicable

## SUPPLEMENTARY MATERIAL

**Supplementary Table 1.** Overall Results

| Sex | Males |  |  |  |  | Females |  |  |  |  |
| --- | --- | --- | --- | --- | --- | --- | --- | --- | --- | --- |
| Treatment | Vehicle | effera™ | effera™ | effera™ | Bovine Lactoferrin | Vehicle | effera™ | effera™ | effera™ | Bovine Lactoferrin |
| Dose Level (mg/kg/day) | 0 | 200 | 600 | 2000 | 2000 | 0 | 200 | 600 | 2000 | 2000 |
| Number of TK Animals | 3 | 6 | 6 | 6 | 6 | 3 | 6 | 6 | 6 | 6 |
| Toxicokinetics<sup>b</sup>: |  |  |  |  |  |  |  |  |  |  |
| C<sub>max</sub> (ng/mL) |  |  |  |  |  |  |  |  |  |  |
| Day 1 | - | 15.2 | 18.8 | 59.1 | 31.2 | - | 851 | 71.2 | 362 | 31.6 |
| Day 28 | - | 1.40 | 85.4 | 75.7 | 29.7 | - | 39.5 | 9.22 | 1530 | 175 |
| AUC<sub>0-24hr</sub> (hr*ng/mL) |  |  |  |  |  |  |  |  |  |  |
| Day 1 | - | NA<sup>c</sup> | 126 | 122 | 21.6 | - | 1270 | 61.3 | 214 | NA<sup>Error!</sup><br>Bookmark not defined. |
| Day 28 | - | NA<sup>c</sup> | 39.5 | 84.1 | NA<sup>c</sup> | - | NA<sup>c</sup> | NA<sup>c</sup> | 511 | NA<sup>c</sup> |
| Anti-LF Antibody |  |  |  |  |  |  |  |  |  |  |
| No. Examined | 3 | 3 | 3 | 3 | 3 | 3 | 3 | 3 | 3 | 3 |
| Anti- effera™ Antibodies |  |  |  |  |  |  |  |  |  |  |
| Treatment | Vehicle | effera™ | effera™ | effera™ | Bovine Lactoferrin | Vehicle | effera™ | effera™ | effera™ | Bovine Lactoferrin |
| Dose Level (mg/kg/day) | 0 | 200 | 600 | 2000 | 2000 | 0 | 200 | 600 | 2000 | 2000 |
| Number of TK Animals | 3 | 6 | 6 | 6 | 6 | 3 | 6 | 6 | 6 | 6 |
| Day 1 | 0 | 0 | 0 | 0 | 0 | 1 | 1 | 0 | 0 | 0 |
| Day 29 | 0 | 2 | 3 | 3 | 0 | 1 | 2 | 1 | 3 | 0 |
| Anti-Bovine Lactoferrin Antibodies |  |  |  |  |  |  |  |  |  |  |
| Day 1 | NA | NA | NA | NA | 0 | NA | NA | NA | NA | 0 |
| Day 29 | NA | NA | NA | NA | 0 | NA | NA | NA | NA | 1 |
| Treatment | Vehicle | effera™ | effera™ | effera™ | Bovine Lactoferrin | Vehicle | effera™ | effera™ | effera™ | Bovine Lactoferrin |
| Dose Level (mg/kg/day) | 0 | 200 | 600 | 2000 | 2000 | 0 | 200 | 600 | 2000 | 2000 |
| Number of Main Study Animals | 10 | 10 | 10 | 10 | 10 | 10 | 10 | 10 | 10 | 10 |
| Noteworthy Findings: |  |  |  |  |  |  |  |  |  |  |
| Died or Sacrificed Moribund | 0 | 0 | 0 | 0 | 0 | 0 | 0 | 0 | 0 | 0 |
| Clinical Observations | - | - | - | - | - | - | - | - | - | - |
| Body Weight | - | - | - | - | - | - | - | - | - | - |
| Body Weight Gains | - | - | - | - | - | - | - | - | - | - |
| Food Consumption<sup>d</sup><br>(g/animal/day) (%) |  |  |  |  |  |  |  |  |  |  |
| Day 1 → 8 | 29.6 | +2 | +2 | -6 | -5 | 20.1 | -2 | -6 | -5 | -9 |
| Day 8 → 15 | 30.1 | +5 | +2 | -4 | -8 | 19.9 | -3 | -9 | +23 | -11 |
| Day 15 → 22 | 30.8 | +3 | 0 | -9 | -8 | 19.7 | +2 | -4 | -6 | -5 |
| Day 22 → 28 | 30.5 | +2 | -1 | -6 | -8 | 20.0 | -7 | -5 | -5 | -4 |
| Ophthalmology | - | - | - | - | - | - | - | - | - | - |
| Hematology | - | - | - | - | - | - | - | - | - | - |
| Coagulation | - | - | - | - | - | - | - | - | - | - |
| Clinical Chemistry | - | - | - | - | - | - | - | - | - | - |
| Urinalysis | - | - | - | - | - | - | - | - | - | - |
| T-cell Dependent Antibody Response | - | - | - | - | - | - | - | - | - | - |
| Treatment | Vehicle | effera™ | effera™ | effera™ | Bovine Lactoferrin | Vehicle | effera™ | effera™ | effera™ | Bovine Lactoferrin |
| Dose Level (mg/kg/day) | 0 | 200 | 600 | 2000 | 2000 | 0 | 200 | 600 | 2000 | 2000 |
| Number of Main Study Animals | 10 | 10 | 10 | 10 | 10 | 10 | 10 | 10 | 10 | 10 |
| Immunophenotyping | - | - | - | - | - | - | - | - | - | - |
| Organ Weights | - | - | - | - | - | - | - | - | - | - |
| Gross Pathology | - | - | - | - | - | - | - | - | - | - |
| Histopathology | - | - | - | - | - | - | - | - | - | - |

GLOSSARY and ENDNOTES
| Abbreviation | Definition |
| --- | --- |
| NA | Not applicable |
| No. | Number |
| TK | Toxicokinetic |
| - | No noteworthy findings |

<sup>a</sup>The positive control article, Keyhole Limpet Hemocyanin, was formulated in sterile phosphate buffered saline, pH 7.4 (minus calcium and magnesium) and Sterile Water for Injection, USP.
<sup>b</sup>Group means are shown. The control group was analyzed for serum concentration; however, group means were not calculated.
<sup>c</sup>AUC not reported due to less than 3 consecutive quantifiable concentrations.
<sup>d</sup> For controls, group means are shown. For treated groups, percent differences from controls are shown. Statistical significance is based on actual data not on the percent differences. All percent difference values are rounded to the nearest whole number.

**Supplementary Table 2.** Analysis of Anti-Recombinant Human Lactoferrin Antibodies in Serum

| Subject Group | Subject | Gender | Day Nominal | Treatment Description | Results |
| --- | --- | --- | --- | --- | --- |
| 6 | 6001 | Male | 1 | Vehicle Control 0 mg/kg | Negative Screen |
| 6 | 6001 | Male | 29 | Vehicle Control 0 mg/kg | Negative Screen |
| 6 | 6002 | Male | 1 | Vehicle Control 0 mg/kg | Negative Screen |
| 6 | 6002 | Male | 29 | Vehicle Control 0 mg/kg | Negative Screen |
| 6 | 6003 | Male | 1 | Vehicle Control 0 mg/kg | Negative Screen |
| 6 | 6003 | Male | 29 | Vehicle Control 0 mg/kg | Negative Screen |
| 6 | 6501 | Female | 1 | Vehicle Control 0 mg/kg | Negative Screen |
| 6 | 6501 | Female | 29 | Vehicle Control 0 mg/kg | Negative Screen |
| 6 | 6502 | Female | 1 | Vehicle Control 0 mg/kg | 225 |
| 6 | 6502 | Female | 29 | Vehicle Control 0 mg/kg | 675 |
| 6 | 6503 | Female | 1 | Vehicle Control 0 mg/kg | Negative Screen |
| 6 | 6503 | Female | 29 | Vehicle Control 0 mg/kg | Negative Screen |
| 7 | 7004 | Male | 1 | effera™ 200 mg/kg | Negative Immunodepletion |
| 7 | 7004 | Male | 29 | effera™ 200 mg/kg | 75.0 |
| 7 | 7005 | Male | 1 | effera™ 200 mg/kg | Negative Screen |
| 7 | 7005 | Male | 29 | effera™ 200 mg/kg | 18200 |
| 7 | 7006 | Male | 1 | effera™ 200 mg/kg | Negative Immunodepletion |
| 7 | 7006 | Male | 29 | effera™ 200 mg/kg | Negative Immunodepletion |
| 7 | 7504 | Female | 1 | effera™ 200 mg/kg | Negative Screen |
| 7 | 7504 | Female | 29 | effera™ 200 mg/kg | 75.0 |
| 7 | 7505 | Female | 1 | effera™ 200 mg/kg | 75.0 |
| 7 | 7505 | Female | 29 | effera™ 200 mg/kg | 18200 |
| 7 | 7506 | Female | 1 | effera™ 200 mg/kg | Negative Immunodepletion |
| 7 | 7506 | Female | 29 | effera™ 200 mg/kg | Negative Screen |
| 8 | 8004 | Male | 1 | effera™ 600 mg/kg | Negative Screen |
| 8 | 8004 | Male | 29 | effera™ 600 mg/kg | 2030 |
| 8 | 8005 | Male | 1 | effera™ 600 mg/kg | Negative Screen |
| 8 | 8005 | Male | 29 | effera™ 600 mg/kg | 675 |
| 8 | 8006 | Male | 1 | effera™ 600 mg/kg | Negative Screen |
| 8 | 8006 | Male | 29 | effera™ 600 mg/kg | 6080 |
| 8 | 8504 | Female | 1 | effera™ 600 mg/kg | Negative Immunodepletion |
| 8 | 8504 | Female | 29 | effera™ 600 mg/kg | 225 |
| 8 | 8505 | Female | 1 | effera™ 600 mg/kg | Negative Screen |
| 8 | 8505 | Female | 29 | effera™ 600 mg/kg | Negative Screen |
| 8 | 8506 | Female | 1 | effera™ 600 mg/kg | Negative Immunodepletion |
| 8 | 8506 | Female | 29 | effera™ 600 mg/kg | Negative Screen |
| 9 | 9004 | Male | 1 | effera™ 2000 mg/kg | Negative Immunodepletion |
| 9 | 9004 | Male | 29 | effera™ 2000 mg/kg | 675 |
| 9 | 9005 | Male | 1 | effera™ 2000 mg/kg | Negative Screen |
| 9 | 9005 | Male | 29 | effera™ 2000 mg/kg | 2030 |
| 9 | 9006 | Male | 1 | effera™ 2000 mg/kg | Negative Screen |
| 9 | 9006 | Male | 29 | effera™ 2000 mg/kg | 18200 |
| 9 | 9504 | Female | 1 | effera™ 2000 mg/kg | Negative Screen |
| 9 | 9504 | Female | 29 | effera™ 2000 mg/kg | 18200 |
| 9 | 9505 | Female | 1 | effera™ 2000 mg/kg | Negative Screen |
| 9 | 9505 | Female | 29 | effera™ 2000 mg/kg | 2030 |
| 9 | 9506 | Female | 1 | effera™ 2000 mg/kg | Negative Screen |
| 9 | 9506 | Female | 29 | effera™ 2000 mg/kg | 2030 |
| 10 | 10004 | Male | 1 | Bovine LF 2000 mg/kg | Negative Immunodepletion |
| 10 | 10004 | Male | 29 | Bovine LF 2000 mg/kg | Negative Immunodepletion |
| 10 | 10005 | Male | 1 | Bovine LF 2000 mg/kg | Negative Screen |
| 10 | 10005 | Male | 29 | Bovine LF 2000 mg/kg | Negative Screen |
| 10 | 10006 | Male | 1 | Bovine LF 2000 mg/kg | Negative Screen |
| 10 | 10006 | Male | 29 | Bovine LF 2000 mg/kg | Negative Screen |
| 10 | 10504 | Female | 1 | Bovine LF 2000 mg/kg | Negative Screen |
| 10 | 10504 | Female | 29 | Bovine LF 2000 mg/kg | Negative Screen |
| 10 | 10505 | Female | 1 | Bovine LF 2000 mg/kg | Negative Immunodepletion |
| 10 | 10505 | Female | 29 | Bovine LF 2000 mg/kg | Negative Screen |
| 10 | 10506 | Female | 1 | Bovine LF 2000 mg/kg | Negative Screen |
| 10 | 10506 | Female | 29 | Bovine LF 2000 mg/kg | Negative Screen |

Abbreviation: LF, lactoferrin

**Supplementary Table 3.** Analysis of Anti-Bovine Lactoferrin Antibodies in Serum

| **Subject Group** | **Subject** | **Gender** | **Day Nominal** | **Treatment Description** | **Result** |
| --- | --- | --- | --- | --- | --- |
| 10 | 10004 | Male | 1 | Bovine LF 2000mg/kg | Negative Immunodepletion |
| 10 | 10004 | Male | 29 | Bovine LF 2000mg/kg | Negative Immunodepletion |
| 10 | 10005 | Male | 1 | Bovine LF 2000mg/kg | Negative Immunodepletion |
| 10 | 10005 | Male | 29 | Bovine LF 2000mg/kg | Negative Immunodepletion |
| 10 | 10006 | Male | 1 | Bovine LF 2000mg/kg | Negative Immunodepletion |
| 10 | 10006 | Male | 29 | Bovine LF 2000mg/kg | Negative Immunodepletion |
| 10 | 10504 | Female | 1 | Bovine LF 2000mg/kg | Negative Immunodepletion |
| 10 | 10504 | Female | 29 | Bovine LF 2000mg/kg | Negative Immunodepletion |
| 10 | 10505 | Female | 1 | Bovine LF 2000mg/kg | Negative Immunodepletion |
| 10 | 10505 | Female | 29 | Bovine LF 2000mg/kg | Negative Immunodepletion |
| 10 | 10506 | Female | 1 | Bovine LF 2000mg/kg | Negative Immunodepletion |
| 10 | 10506 | Female | 29 | Bovine LF 2000mg/kg | 150 |

Abbreviation: LF, lactoferrin

**Supplementary Figure 1.**
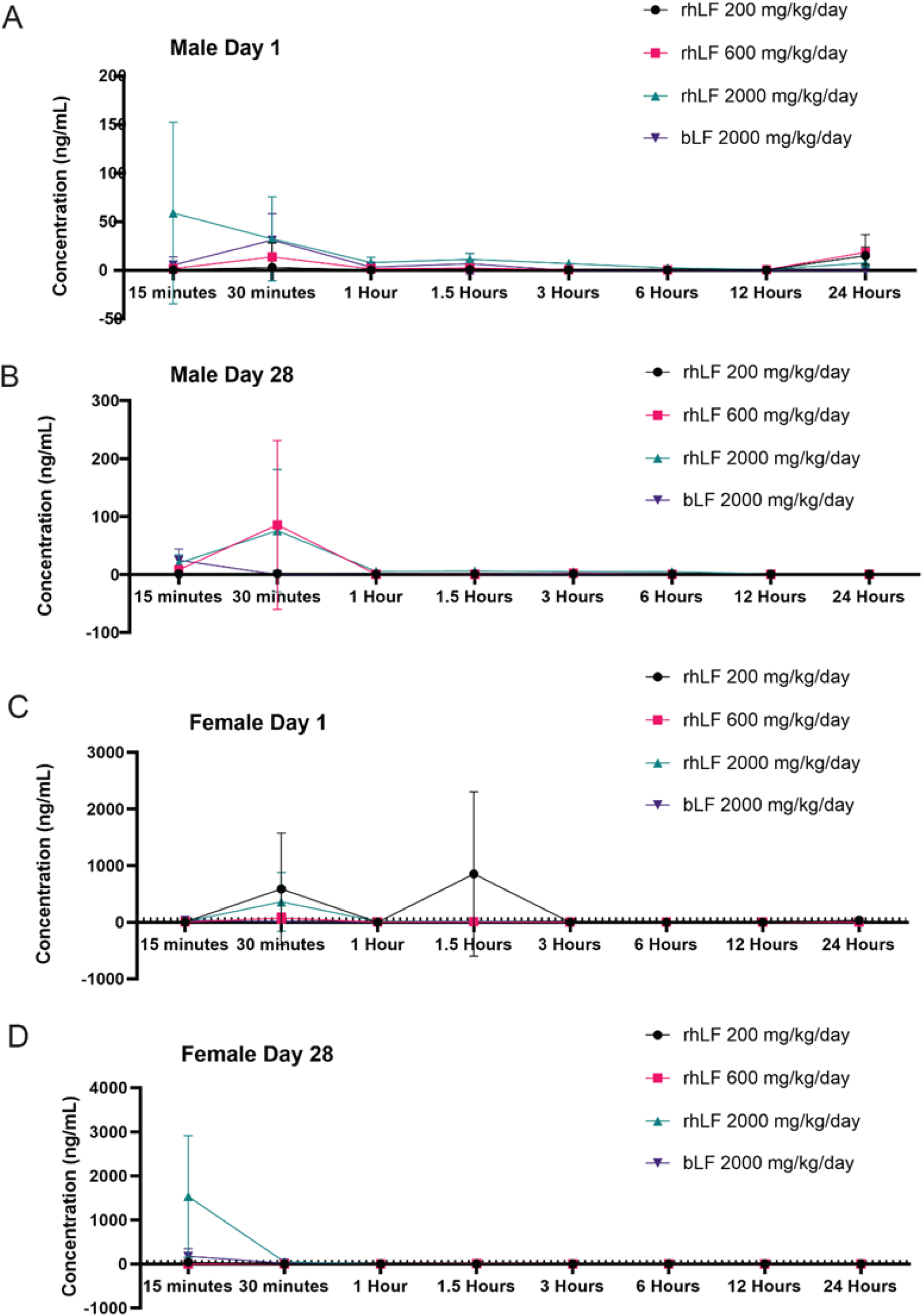
Postprandial levels of LF detected in the serum. Serum was collected over multiple time points after administration on Day 1 and Day 28 of dosing. ECLIA was performed to detect rhLF and ELISA was performed to detect bLF in the serum. A. Day 1 LF results in male rats B. Day 28 LF results in male rats. C. Day 1 LF results in female rats. D. Day 28 results in female rats. N = 3 animals per time point. rhLF = Effera^TM^, bLF = bovine LF

